# Bayesian inference of ancestral host-parasite interactions under a phylogenetic model of host repertoire evolution

**DOI:** 10.1101/675017

**Authors:** Mariana P Braga, Michael Landis, Sören Nylin, Niklas Janz, Fredrik Ronquist

## Abstract

Intimate ecological interactions, such as those between parasites and their hosts, may persist over long time spans, coupling the evolutionary histories of the lineages involved. Most methods that reconstruct the coevolutionary history of such associations make the simplifying assumption that parasites have a single host. Many methods also focus on congruence between host and parasite phylogenies, using cospeciation as the null model. However, there is an increasing body of evidence suggesting that the host ranges of parasites are more complex: that host ranges often include more than one host and evolve via gains and losses of hosts rather than through cospeciation alone. Here, we develop a Bayesian approach for inferring coevolutionary history based on a model accommodating these complexities. Specifically, a parasite is assumed to have a host repertoire, which includes both potential hosts and one or more actual hosts. Over time, potential hosts can be added or lost, and potential hosts can develop into actual hosts or vice versa. Thus, host colonization is modeled as a two-step process, which may potentially be influenced by host relatedness or host traits. We first explore the statistical behavior of our model by simulating evolution of host-parasite interactions under a range of parameters. We then use our approach, implemented in the program RevBayes, to infer the coevolutionary history between 34 Nymphalini butterfly species and 25 angiosperm families.

---

Extant ecological interactions, such as those between parasites and hosts, are often the result of a long history of coevolution between the involved lineages (Elton 1946; Klassen 1992).Specialization is predominant among parasites (including parasitic herbivorous insects; Forister et al. 2015), but host associations are not static: they continuously evolve over time via gains and losses of hosts (Janz and Nylin 2008; Nylin et al. 2018). The colonization of new hosts and loss of old hosts not only shape the evolutionary trajectories of the interacting lineages, but can also have large effects at ecological timescales (Nosil 2002; Calatayud et al. 2016). These effects are evident, for example, with emerging infectious diseases and zoonotic diseases (Acha and Szyfres 2003), which involve colonization of new hosts within and among groups of domesticated species (Subbarao et al. 1998), wildlife (Fisher et al. 2009), and humans (Hahn et al. 2000). Unraveling the processes underlying changes in species associations is thus key to understanding evolutionary and ecological phenomena at various timescales, such as the emergence of infectious diseases, community assembly, and parasite diversification (Hoberg and Brooks 2015).

Many methods developed to study historical associations focus on congruence between host and parasite phylogenies (Brooks 1979; Huelsenbeck et al. 1997; de Vienne et al. 2013). Such methods largely fall into two main classes of cophylogenetic approaches: (1) topology- and distance-based methods, which estimate the congruence between two phylogenies (Legendre et al. 2002), and (2) event-based methods, which map the parasite phylogeny onto the host phylogeny using evolutionary events (Ronquist 2003). Typically, cospeciation is the null hypothesis in these methods, where host shifts are invoked only to explain deviations from cospeciation (de Vienne et al. 2013). Moreover, most of these methods do not allow ancestral parasites to be associated with more than one host lineage, thus failing to account for a potentially important driver of parasite diversification (Janz and Nylin 2008).

An alternative approach to studying coevolving host-parasite associations is to perform ancestral state reconstructions of individual host taxa onto the parasite phylogeny and combine the ancestral host states a posteriori into inferred host ranges (e.g. Nylin et al. 2014). Even though this approach allows ancestral parasites to have multiple hosts, it assumes that the associations between the parasite and each host evolve independently. This has a number of serious drawbacks. For instance, ancestral parasites may be inferred to have an unrealistically high number of hosts, or no host at all. Furthermore, the more narrowly circumscribed the host taxa are, the more likely it is that ancestral parasite lineages are reconstructed as having no hosts. In addition, the independence assumption causes the phylogenetic relationships among hosts to be ignored, meaning that the model assigns equal rates to all colonizations of new hosts regardless of how closely related the new host is to the current hosts being used by the parasite.

A desirable model of host usage should therefore allow parasites to have multiple hosts, while also allowing for among-host (or context-dependent) effects to influence ancestral host use estimates and gain and loss rates in whatever manner explains the biological data best. One possible solution is to restate the problem of host-parasite co-evolution in terms of historical biogeography. For instance, the Dispersal-Extirpation-Cladogenesis (DEC) model of Ree et al. (2005) allows species ranges to stochastically evolve as a set of discrete areas over time through area gain events (dispersal), area loss events (extirpation), and cladogenetic events (range inheritance patterns that reflect speciational models). Although these methods are designed for biogeographic inference, a similar approach is clearly suitable for more realistic modeling of host-parasite coevolution dynamics, where colonization and loss of hosts (instead of discrete areas) is modeled as a continuous-time Markov process (e.g. Hardy 2017). In biogeography, the colonization of a new area or the disappearance from a previously occupied area is modeled as a binary trait: the species is either present or absent in the area. While this binary view might be simple but useful in biogeography, it may be too simplistic for use in the coevolution between hosts and parasites. For instance, it is known that butterflies can utilize a range of plants that they do not regularly feed on in the wild, and it has been suggested that these potential hosts have played an important role in the evolution of host use in butterflies, by increasing the variability in host use through time and across clades (Janz et al. 2016; Braga et al. 2018). This hypothesis can only be directly tested, however, if we explicitly model the evolution of host use as a two-step process, which cannot be done with the binary methods that are used today to study host-parasite coevolution or biogeography.

Here, we propose a model where a parasite is assumed to have a host repertoire, defined as the set of all potential and actual hosts for that parasite. In this model, the colonization of a new host involves two steps: first, the parasite gains the ability to use the new host (it becomes a potential host), and then starts actually using it in nature (it becomes an actual host). These two steps can be interpreted as the inclusion of the new host into the fundamental and then into the realized host repertoire of the parasite - analogous to fundamental and realized niche (Nylin et al. 2018; Larose et al. 2019). Similarly, the complete loss of a host from a parasite’s realized repertoire involves two steps. First, it changes from an actual to a potential host, and then it is lost completely from the host repertoire. For example, if the geographic range of a host contracted to become allopatric with respect to a parasite’s geographic range, the host would remain as part of the fundamental repertoire until the parasite completely lost the ability to use the host, in which case the host would be lost from the repertoire. Even when in sympatry, the evolution of a new defense mechanism by the host may prevent the parasite from using that host. However, since host use is a complex and multidimensional trait, it is unlikely that a parasite loses all the machinery necessary to use a host in one single event, and it may well retain some ability to survive on the host. Thus, three host-parasite association states are necessary for such a two-step model: the host is used (actual host), the parasite has some ability to use the host but does not use it in nature (potential host), and the parasite cannot use the host (non-host).

In this paper, we develop a Bayesian approach to coevolutionary inference based on such a model of host repertoire evolution, inspired by the previous work on similar biogeographic inference problems by Landis et al. (2013). The basic binary biogeographic model, when applied to coevolution, accommodates both multiple ancestral hosts and changes in host configurations over time that correspond to evolutionary changes in host lineages or host traits. We extend this model to also include a two-step host colonization process, such that the fundamental host repertoire can persist over time and affect the evolution of the realized repertoire. We have implemented the model in RevBayes (Höhna et al. 2016), allowing us to perform simulation as well as Bayesian Markov chain Monte Carlo (MCMC) inference under the model. This Bayesian framework allows one to estimate the joint distribution of host gain and loss rates, the effect (if any) of phylogenetic distances among hosts upon host gain rates, and the historical sequences of evolving host repertoires among the parasites. Using simulations, we explore the statistical behavior of our approach, and demonstrate its empirical application with an analysis of the coevolution between Nymphalini butterflies and their angiosperm hosts.

## Methods

### Model description

We are interested in modeling the evolution of ecological interactions between M extant parasite taxa and N host taxa, where each parasite uses one or more hosts. Rooted and time-calibrated phylogenetic trees describe the evolutionary relationships among the M parasite taxa and among the N host taxa. In this study, the trees are considered to be known without error. In principle, it would be straightforward for the model to accommodate phylogenetic uncertainty in the host or parasite trees but MCMC inference may prove challenging under such conditions.

Each parasite taxon has a host repertoire, which is represented by a vector of length N that contains the information about which hosts the given parasite uses. The interaction between the m-th parasite and the n-th host is denoted x_m,n_. At any given time, each host taxon can assume one of three states with respect to a parasite lineage: x_m,n_ is equal to 0 (non-host), 1,m (potential host), or 2 (actual host). Criteria for how to code non-host, potential host, and actual host states will depend on the host-parasite system under study; below, we provide criteria for our Nymphalini dataset that may act as guidelines. We allow all host repertoires in which the parasite has at least one actual host. Thus, the state space, S, includes 3^*N*^ – 2^*N*^ host repertoires for *N* hosts.

Here we define the transition from state 0 to state 1 as the gain of the ability to use the host, and the transition from state 1 to state 2 as the time when the parasite actually starts to use the host in nature. If we assume that gains and losses of hosts occur according to a continuous-time Markov chain, the probability of a given history of association between a parasite clade and their hosts can be easily calculated (Ree and Smith 2008). This calculation is based on a matrix, **Q**, containing the instantaneous rates of change between all pairs of host repertoires, and thus describing the Markov chain. Based on the **Q** matrix, it is possible to calculate the transition probability of the observed host repertoires at the tips of the parasite tree by marginalizing over the infinite number of histories that could produce the observed host repertoires. Unfortunately, computing these transition probabilities becomes intractable as the number of host repertoire configurations, *S*, grows large. Modeling host repertoire evolution for host repertoire size *N* = 7 requires an *S* × *S* rate matrix defined for *S* = 3^7^ *–* 2^7^ = 2059, causing **Q** to be too large for efficient inference. In order to handle large host repertoires, we numerically integrate over possible histories using data augmentation and MCMC rather than analytically computing the probabilities using matrix exponentiation. This data augmentation approach has been used to model sequence evolution for protein-coding genes (Robinson 2003) and historical biogeography (Landis et al. 2013; Quintero and Landis 2019), suggesting the framework may be useful to model host-parasite interactions as well. In this study, we assume that both daughter lineages identically inherits their host repertoires from their immediate ancestor at the time of cladogenesis.

We define a model where the gain of a host (both 0→1 and 1→2) depends on the phylogenetic distance between the available hosts and those currently used by a lineage. Figure 1 schematically illustrates the evolutionary dynamics of the model using *M* = 4 parasite species and *N* = 5 host species, while assuming that host gain rates are independent (Fig. 1a,c) or dependent (Fig. 1b,d) of phylogenetic distances among hosts. To formalize these dynamics, let 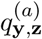 be the rate of change from host repertoire **y** to repertoire **z** by changing the state of host *a*. Also, let *λ*_*ij*_ be the rate at which an individual host changes from state *i* to state *j*, and *η*(**y**, *a, β*) be a phylogenetic-distance rate modifier. The phylogenetic-distance rate modifier function, *η*, rescales the base rate of host gain to allow new hosts that are closely related to the parasite’s current hosts to be colonized at higher rates than distantly related hosts. We define the instantaneous rate of change as

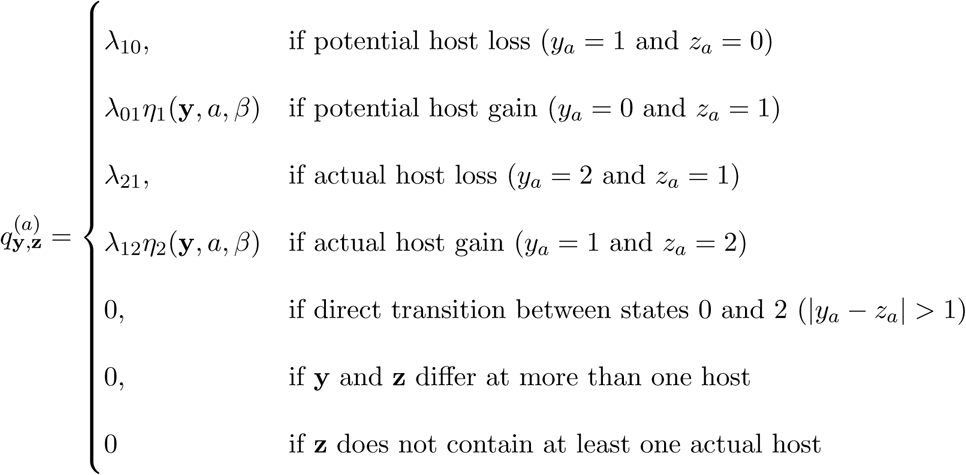

and the phylogenetic-distance rate modifier function as

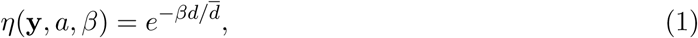

where *β* controls the effect of *d*, the average pairwise phylogenetic distance between the new host, *a*, and the hosts currently occupied in **y**; and 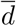 is the average phylogenetic distance between all pairs of hosts. Pairwise phylogenetic distance is defined as the sum of branch lengths separating two leaf nodes. The difference between *η*_1_ and *η*_2_ is that in the first, pairwise distances are calculated between the new host and all potential and actual hosts, while in the second only actual hosts are included. This allows for a model formulation where the effect of host distances on *λ*_01_ and on *λ*_12_ are independent, while still allowing a formulation where they are equal. If *β* = 0, the gain rate of host *a* is equal to the unmodified gain rate, *λ*_01_ or *λ*_12_. If *β >* 0, the gain rate of phylogenetically close hosts is higher than distant hosts.

**Figure 1:**
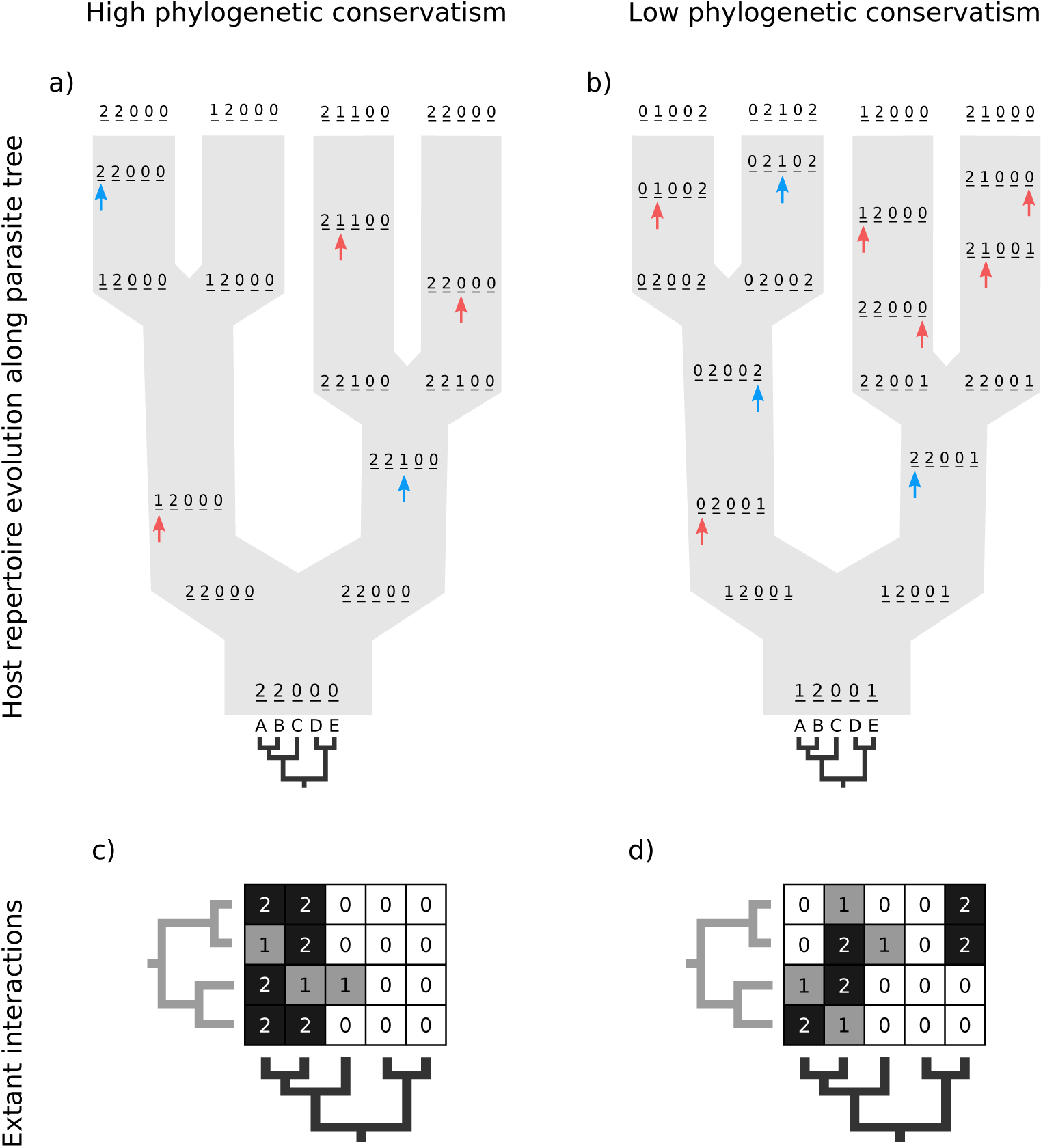
Host repertoire evolution along a hypothetical tree and resulting host-parasite interactions. Two examples of coevolutionary histories between four parasites and five hosts are shown to illustrate how the model works. Host repertoires evolve by gains (0→1 and 1→2, blue arrows) and losses (1→0 and 2→1, red arrows). Coevolutionary histories in **a** and **b** produce the interactions in **c** and **d** respectively. In **c** and **d**, each column represents one host and each row represents the host repertoire of one parasite. High phylogenetic conservatism is produced when the rate of repertoire evolution, *µ*, is low and the effect of the phylogenetic distance between hosts, *β*, is high. Conversely, low phylogenetic conservatism is produced when *µ* is high and *β* is low.

We fit this model using the Bayesian data augmentation strategy described in Landis et al. (2013). The method estimates the joint posterior probability of model parameters, *θ* = (*µ, λ, β*), and data-augmented evolutionary histories, *X*_aug_, conditional on the observed host repertoire data, *X*_obs_, and the parasite phylogeny, Ψ_p_, and the host phylogeny, Ψ_h_, using MCMC. To sample values from the posterior, *P* (*X*_aug_, *θ | X*_obs_, Ψ_*p*_, Ψ_*h*_), new parameter values for *µ, λ*, and *β* are proposed using standard Metropolis-Hastings proposals for updating simple parameters (Hastings 1970). Analogously, our MCMC stochastically proposes and/or accepts new augmented host repertoire histories using the Metropolis-Hastings algorithm. Augmented histories are proposed using two types of MCMC moves: branch-specific moves and node-and-branch moves. Branch-specific moves propose a new augmented history by sampling a branch from the phylogeny uniformly at random, then proposing new histories for a subset of host-characters using the rejection sampling method of Nielsen (2002) under the assumption that all host characters evolved under mutual independence (β = 0); this assumption allows us to rapidly propose new augmented histories. Although augmented histories are proposed assuming host characters evolve independently, we compute the acceptance probability for the branch-specific move by considering the full-featured model probability that allows for non-independent rates of character change when calculating the Metropolis-Hastings ratio. Thus, the augmented histories are sampled in proportion to their posterior probabilities under the full model. Node-and-branch moves involves sampling new host repertoire states for a node sampled uniformly at random within the parasite tree, along with the three branches incident to the node. Together, the branch-specific moves, the node-and-branch moves, and the parameter moves allow MCMC to estimate the posterior probability of combinations of host repertoire histories and evolutionary parameters. Further details are provided in Landis et al. (2013).

### Model selection

When *β* = 0, the phylogenetic-distance dependent model, *M*_*D*_ becomes a mutual-independence model, *M*_*0*_, where the interaction between the parasite and each host evolves independently.

These models are therefore nested (*M*_0_ *⊆ M*_*D*_) and we can compute Bayes factors for model *M*_*D*_ over model *M*_0_ using the Savage-Dickey ratio (Verdinelli and Wasserman 1995; Suchard et al. 2001), defined as

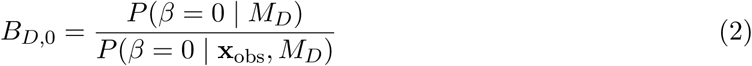

where *P*_*D*_(*β* = 0 *| M*_*D*_) is the prior probability and *P* (*β* = 0 *|* **x**_obs_, *M*_*D*_) is the posterior probability, both defined in terms of the phylogenetic-distance dependent model, *M*_*D*_, at the restriction point *β* = 0 where *M*_*D*_ and *M*_*0*_ are equivalent. While we could directly compute the prior probability of *β* = 0, we approximated the posterior at *β* = 0 using a kernel density estimator with a gamma function, which only takes positive values, and a bandwidth of 0.02. To interpret if and how Bayes factors favored the phylogenetic-distance dependent model, *M*_*D*_, we followed the guidelines of Jeffreys (1961): model *M*_0_ is favored for Bayes factors with values less than 1, insubstantial support is awarded to model *M*_*D*_ for values between 1 and 3, substantial support for values between 3 and 10, strong support for values between 10 and 30, very strong support for values between 30 and 100, and decisive support for values greater than 100.

### Data analysis

#### Simulation study

We simulated 50 datasets for each of nine combinations of values for the rate of host-repertoire evolution, *µ* (0.01, 0.04, and 0.1), and values of *β* (0, 1, and 4). These parameter combinations produce datasets with varying degrees of phylogenetic conservatism for both parasites and hosts (Fig. 2). Each dataset contained 34 insects and 25 hosts, and was produced by simulating host repertoire evolution in the parasite tree used in the empirical study (see below). Host gain and loss rates were chosen to resemble the rates inferred from the empirical analysis. This simulation was designed to assess our statistical power to detect the effect of phylogenetic distance among hosts upon host gain rates given the size of our empirical dataset and the type of variation we expected it to contain.

**Figure 2:**
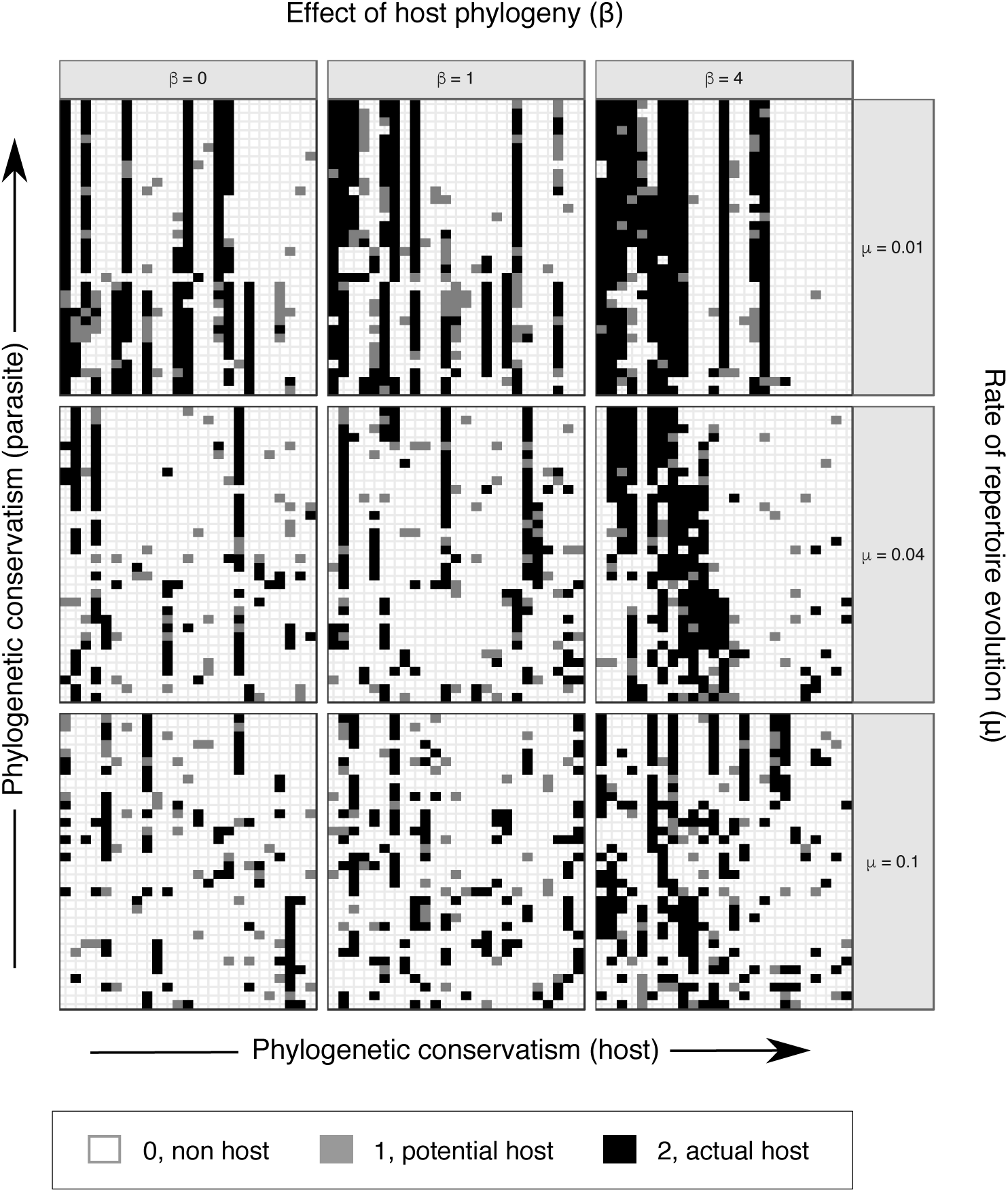
Simulated datasets for nine parameter combinations. Interactions between Nymphalini butterflies and their host plants for one of 50 simulations with each parameter combination. In each of the nine datasets, each column represents one host in the repertoire and each row shows the host repertoire of one butterfly species. When phylogenetic conservatism in host-parasite interactions is low for both hosts and parasites, the interactions are more randomly spread (matrix at bottom-left corner). As phylogenetic conservatism among parasites increases, host repertoires (rows) become more similar (upper matrices). When there is phylogenetic conservatism among hosts, host repertoires include more closely-related hosts (neighbouring columns; matrices to the right)

We ran independent MCMC analyses for each set of 50 datasets, under the phylogenetic-distance dependent model. We then quantified how well the posterior probabilities of coevolutionary histories correspond to the true history known from each simulation. Specifically, we first computed the posterior probability of interaction between each host and each internal node in the butterfly tree, for states 1 and 2 separately. Then, we calculated the sum of squared differences between each posterior probability (0 ≤ *P* ≤ 1) and the corresponding truth for that simulation (1, if the host was on the given state in the simulated dataset; 0, if not). This error term increases as the inferred ancestral host repertoires become less accurate.

#### Empirical study

In order to validate our method, we compiled data from the literature for butterflies from the tribe Nymphalini (Nymphalidae) and their host plants (see Supplementary Information for reference list). We chose this butterfly clade because we expect that a large fraction of the real potential hosts are known, as there has been systematic experimental studies of larval feeding ability. The dataset included 34 butterflies species and plants from 16 angiosperm families (Figs. S1 and S2). For each butterfly species, host plants commonly used in nature were coded as ‘actual hosts’ and plants never used were coded as ‘non-hosts’. Plants that are not commonly used in nature, but for which there is strong evidence (field observation or experiment) that the larvae can feed upon them, were coded as ‘potential hosts’.

Because we lack the information on potential hosts for most host-parasite systems (i.e. hosts are usually only classified as hosts or non-hosts), we tested whether our model is able to recover the same parameter estimates and coevolutionary histories when all the potential hosts are coded as non-hosts. For that, we ran the same analysis as for the full dataset, but first removed all the 1s from the empirical dataset. Then we compared the posterior probabilities inferred from the two datasets. To assess the similarities between the coevolutionary histories inferred using the different datasets, we calculated summary statistics for the absolute difference in probability of each interaction between hosts and internal nodes in the butterfly tree.

For both the simulation and empirical studies we used the phylogenetic relationships between butterfly species in the Nymphalini tribe as proposed by Chazot (unpublished, Fig. S3) and the phylogenetic relationships between angiosperm families proposed by Magallón et al. (2015). Although our framework allows the inclusion of a large number of hosts in the same analysis, computational time increases significantly with the size of the host repertoire. We therefore chose to include 25 hosts, which allows the inclusion of all host lineages used by any of the butterflies. To ensure the inclusion of all plant lineages that might have been used as hosts in the past, we pruned the angiosperm phylogenetic tree so that all 16 families in the dataset were included, and the remaining branches were collapsed to more ancestral nodes until only 25 tips were left. We then pruned all the branches leading up to the tips to the time of origin of the butterfly clade (approx. 22 Ma), and this pruned tree was then used to calculate phylogenetic distances between hosts. To simplify the analysis, we hold the phylogenetic distances between plant families constant, independent of geological time, even though the distances would be expected to increase as evolution proceeds towards the recent.

We summarized inferred coevolutionary histories in two ways. First, we calculated the posterior probability for fundamental and realized host repertoires at internal nodes of the Nymphalini phylogeny based on the frequency with which states 1 and 2 were sampled for each host during MCMC. Second, in order to reduce the dimensionality of the host repertoire and facilitate visualization of ancestral state reconstructions, we assigned hosts to modules based on extant butterfly-plant interactions (Fig. S2). Modules are groups of plants and butterflies that interact more with each other than with other taxa, thus host plants are assigned to the same module when they are used by the same butterflies. To identify the modules, we used a simulated annealing algorithm that maximizes the index of modularity. Specifically, we used Newman and Girvans metric (Newman and Girvan 2004) modified for bipartite networks (Barber 2007) as implemented in the software MODULAR (Marquitti et al. 2014).

#### Software configuration

All analyses were performed in RevBayes (Höhna et al. 2016). For the simulated data, we ran two independent MCMC analyses for 10^5^ cycles, sampling parameters and node histories every 50 cycles, and discarding the first 5 × 10^4^ as burnin. For the empirical data, we ran five independent MCMC analyses, each set to run for 10^6^ cycles, sampling every 50 cycles, and discarding the first 10^5^ as burnin. To verify that MCMC analyses converged to the same posterior distribution, we applied the Gelman diagnostic (Gelman and Rubin 1992) provided through the R package coda (Plummer et al. 2006). For both simulated and empirical datasets, we used the following priors: β ∼Exponential(1), µ ∼ Exponential(10), andλ ∼ Dirichlet(1, 1, 1, 1). Analysis scripts and data files are available at https://github.com/mpiresbr/host_repertoire. A RevBayes tutorial for the empirical analysis will be soon available at https://revbayes.github.io/tutorials#host_rep.

## Results

### Simulation study

Posterior distributions of parameter values for the 9×100 MCMC analyses are shown in Figure 3. Overall, the model was able to accurately recover the true simulation parameters (true value within 95% highest posterior density, or HPD). However, accuracy decreased with increasing rate of host repertoire evolution, possibly due to character saturation.

We performed model selection based on Bayes factors. Considering that the prior distribution is *β* ∼ *Exponential*(1), a high marginal posterior probability for *β* = 0 under *M*_*D*_ is necessary to result in a Bayes factor *<* 1 and thus selection of *M*_0_. For simulations with *β* = 0, the correct model, *M*_0_, was selected in more than 60% of the simulations, and most of the remaining simulations gave insubstantial support to *M*_*D*_ (Fig. 4). When *β* = 1, Bayes factors correctly selected *M*_*D*_ in the majority of cases, but strong support for *M*_*D*_ was only achieved in simulations with *β* = 4, particularly when the rate of evolution was highest (*µ* = 0.1).

We then compared the true coevolutionary history of each simulation to the corresponding posterior distribution of the sampled coevolutionary histories (Fig. 5). The estimation error, that is, the sum of squared differences between estimated and true coevolutionary histories, was very low when the rate of host-repertoire evolution was lowest (*µ* = 0.01), but also when the phylogenetic-distance power was highest (*β* = 4). This means that accuracy in the estimation of coevolutionary history increases with phylogenetic conservatism on both the butterfly and the plant trees. Overall, error was higher on the estimation of actual hosts (state 2) than potential hosts (state 1).

### Empirical study

The estimated mean rate of host repertoire evolution for Nymphalini was *µ* = 0.025, the mean phylogenetic-distance power was *β* = 0.51, and the mean gain/loss rates were *λ*_01_ = 0.012, *λ*_10_ = 0.6, *λ*_12_ = 0.27, and *λ*_21_ = 0.12 (Fig. 6, blue). Our method recovered similar parameter estimates for the empirical dataset when omitting the intermediate state at the tips – i.e. coding all potential hosts (state 1) as non-hosts (state 0): *µ* = 0.031, *β* = 0.39, *λ*_01_ = 0.001, *λ*_10_ = 0.71, *λ*_12_ = 0.28, and *λ*_21_ = 0.01 (Fig. 6, orange). The posterior distributions from analyses with and without the intermediate state at the tips diverged the most for the rate parameters associated with the transition to the intermediate state, *λ*_01_ and *λ*_21_. In both cases the transition rate was underestimated when 1s were removed from the dataset. Bayes factors selected the independence model, *M*_0_, for both the full dataset (BF = 0.43) and when 1s were removed from tip states (BF = 0.40).

Finally, we reconstructed the fundamental and realized host repertoires at internal nodes of the Nymphalini phylogeny based on the sampled histories during MCMC. Coevolutionary histories inferred using the datasets with and without potential hosts were very similar, with mean difference in interaction probability of 0.003. Thus, we only show the ancestral states inferred from the full, three-state dataset (Figs. 7 and S4). To facilitate visualization of the ancestral state reconstruction, we grouped the 16 parasitized host families into five modules, as identified by the simulated annealing algorithm (Fig. S2). Nine families (representing three modules) were inferred to be used by ancestral Nymphalini species with high probability.

We found strong support for the association between the ancestor of all Nymphalini butterflies and Urticaceae hosts (and Cannabaceae to a lesser degree, Fig. S4). All other host families have been colonized in the last 15 Myr, after the divergence of the two largest clades within Nymphalini, *Vanessa* and *Nymphalis* + *Polygonia*. Most species within *Vanessa*, both extant and ancestral, are specialists on Urticaceae. *V. virginiensis* and *V. cardui* are the only extant species that use more than two host families, and these hosts have likely been colonized by their most recent common ancestor (node 38 in Fig. 8). On the other hand, the variation in host use in the *Nymphalis* + *Polygonia* clade seems to be the result of host colonizations by multiple species along the diversification of the clade. For example, in Fig. 8 we can see the colonization of potential hosts by the ancestor of *P. c-album* and *P. faunus* (node 53) as well as strong specialization on a new host by *Kaniska canace*.

## Discussion

The method we develop here to infer the evolutionary history of host-parasite associations has many advantages over previous approaches. First, it is based on stochastic models and on established principles of statistical inference, which means that it provides a robust framework for characterizing the evolutionary processes that shape host-parasite associations and for selecting among alternative coevolutionary models. Second, our model introduces the novel concept of a host repertoire, which we think is an important step forward. Besides accounting for the possibility of parasites having more than one host over time scales of macroevolutionary significance, we can now directly infer the influence of host relatedness and host traits on the process of gaining new hosts. Third, the stochastic model of host-parasite coevolution that we introduce here is, to our knowledge, the first that explicitly accounts for evolution of the fundamental host repertoire. By recognizing the fact that a parasite may have potential hosts in addition to its actual hosts, and that the set of potential hosts may persist over time, the dynamic of the model changes. What would otherwise have appeared as remarkable repeated patterns of colonization of the same host lineages can now be explained as the effect of frequent transitions between potential and actual hosts in an otherwise conserved host repertoire.

Our model can readily be extended in many interesting ways. The version we present here accounts for the effect of host phylogeny by allowing the rate of host gain to depend on host relatedness. For simplicity, we assumed that the number of available hosts and host relatedness remain constant over geological time. This would be appropriate for a group of parasites that radiated after the relevant host lineages had been formed, which is arguably the case for the empirical example we chose. However, it should be relatively straightforward to extend our framework to account for more complex dependencies on host phylogeny. For instance, the host configurations could be modeled as changing over time, reflecting host cladogenesis.

Another interesting direction for future research would be to modify the particular ways in which hosts and parasites coevolve. We note, for example, that Fig. 7 shows that host repertoires of *Vanessa* species overlap very little with the host repertoires of *Nymphalis* + *Polygonia* species, but it is not immediately clear what drives this pattern. One could design a model that allows the rates of host gain and loss to be influenced by evolving host traits — like secondary metabolites, growth form, or phenology, to mention a few examples relevant for insect-host plant associations — in addition to relatedness among hosts. Or, one might extend the model to allow closely related parasite lineages to competitively exclude one another from host usage, similar to how competing lineages might exclude one another from geographical regions (Quintero and Landis 2019). Finally, one might introduce a biogeographical component to the coevolutionary process, requiring parasites to be in sympatry with their actual hosts, while allowing parasites to be in sympatry or allopatry with their potential hosts. Statistically comparing such model variants will help illuminate drivers of host-parasite co-evolution.

A potential concern with our approach is that already the basic version of the model is fairly parameter-rich. Given the type and amount of data that we can likely collect on host-parasite associations, is there enough statistical power to select among the models of interest? And is it possible to infer the model parameters of interest with a reasonable degree of accuracy?

Overall, our results are encouraging in this respect. The simulations indicate that it is possible to infer the true parameter values of the basic model regardless of the level of phylogenetic conservatism in both parasites and hosts (Fig. 3). When the rates of colonization of new hosts are strongly dependent on the phylogenetic relatedness of hosts, then we are also able to distinguish between models with or without host relatedness effects using Bayes factors (Fig. 4). However, our ability to select the correct model decreases when the effect of host phylogenetic relatedness is low (*β* ≤ 1), that is, when models become more similar. Further studies will have to show to what extent the sensitivity of the model test can be increased by selecting appropriate priors and improving the sampling of parameter space close to the boundary condition satisfying the restricted model. One option is to relax the assumption that *β* is non-negative, which would simplify the sampling of values close to *β* = 0. It will also be important to explore how dataset sizes and tree shapes, for both hosts and parasites, influence our ability to distinguish the models when the effect of host phylogeny is small.

**Figure 3:**
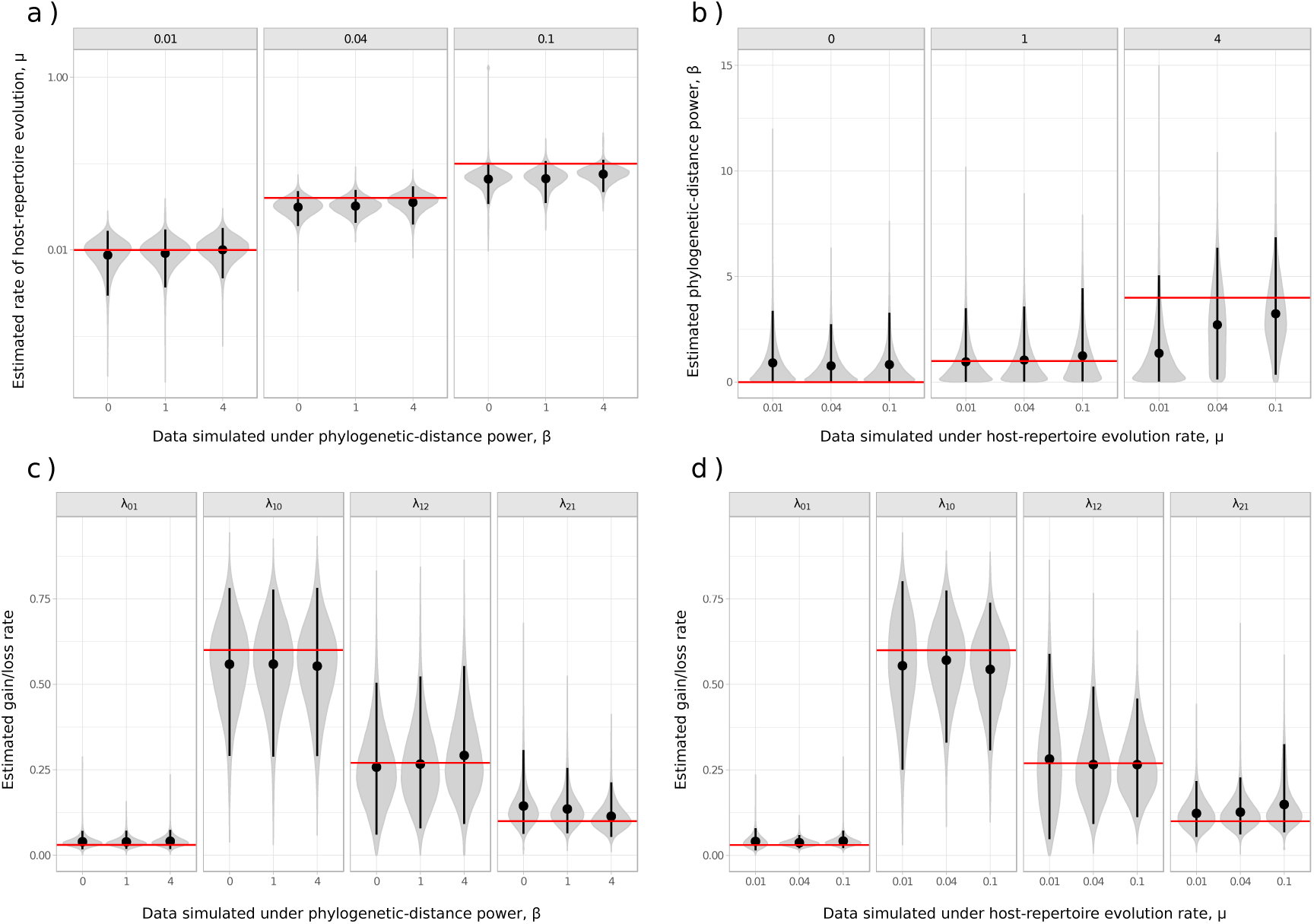
Posterior densities of parameters in the simulation study. Panels **a** and **b** are faceted by true parameter values of *µ* and *β*, respectively. Fifty datasets were simulated for each combination of *β ∈{*0, 1, 4*}* and *µ ∈{*0.01, 0.04, 0.1*}*, while *λ*_01_ = 0.03, *λ*_10_ = 0.6, *λ*_12_ = 0.27, and *λ*_21_ = 0.1 were held constant. For each parameter combination, the posterior distributions of the two MCMC samples of the 50 datasets were combined. Means are represented by black dots, black vertical lines show the 95% HPD, and red horizontal lines mark the true parameter value used in the simulations. Y-axis in panel **a** is in *log*_10_ scale for better visualization.

**Figure 4:**
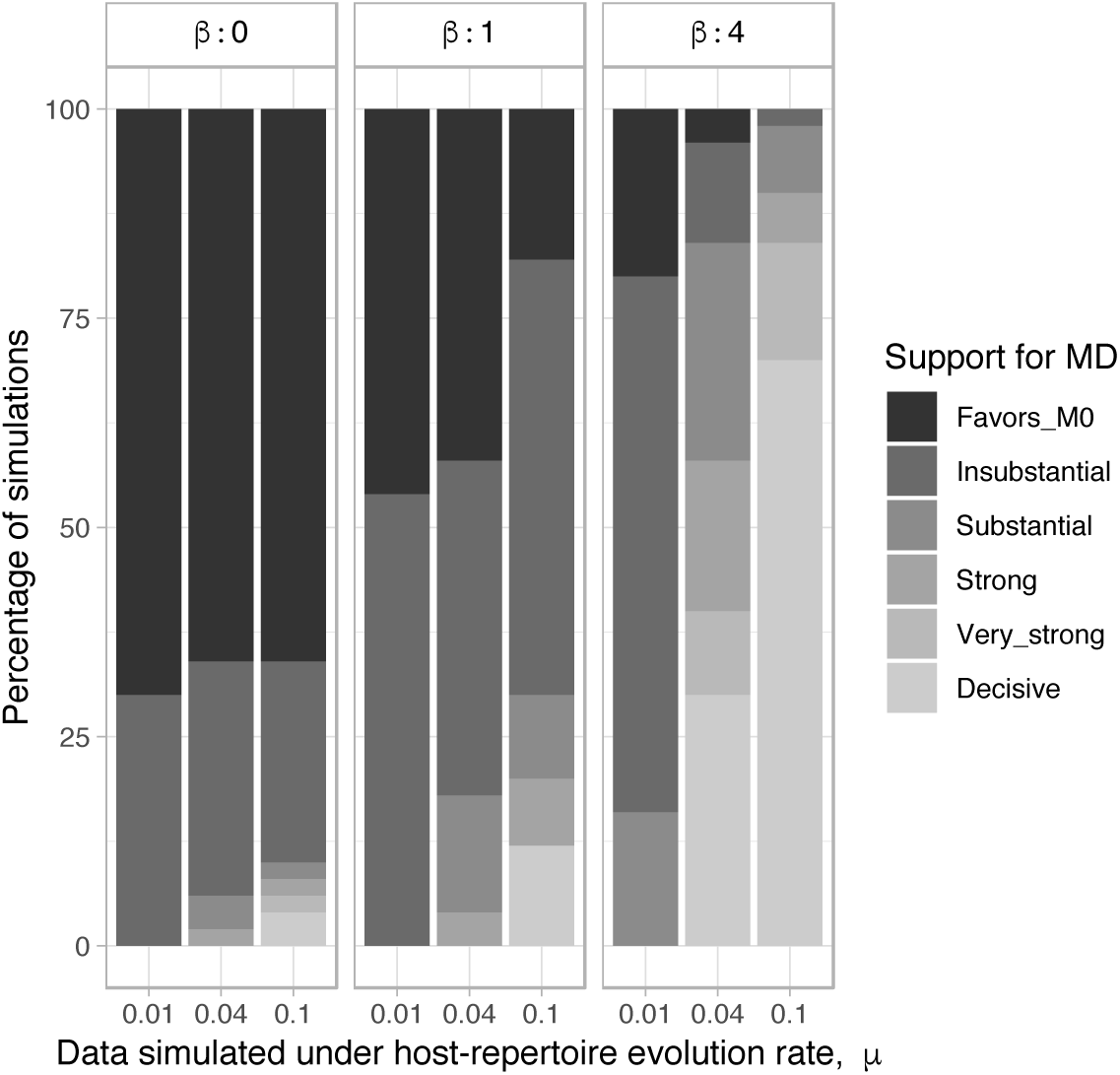
Distribution of Bayes factors for the simulation study. Each column corresponds to the strength of support per 2 ×50 MCMC analyses.

**Figure 5:**
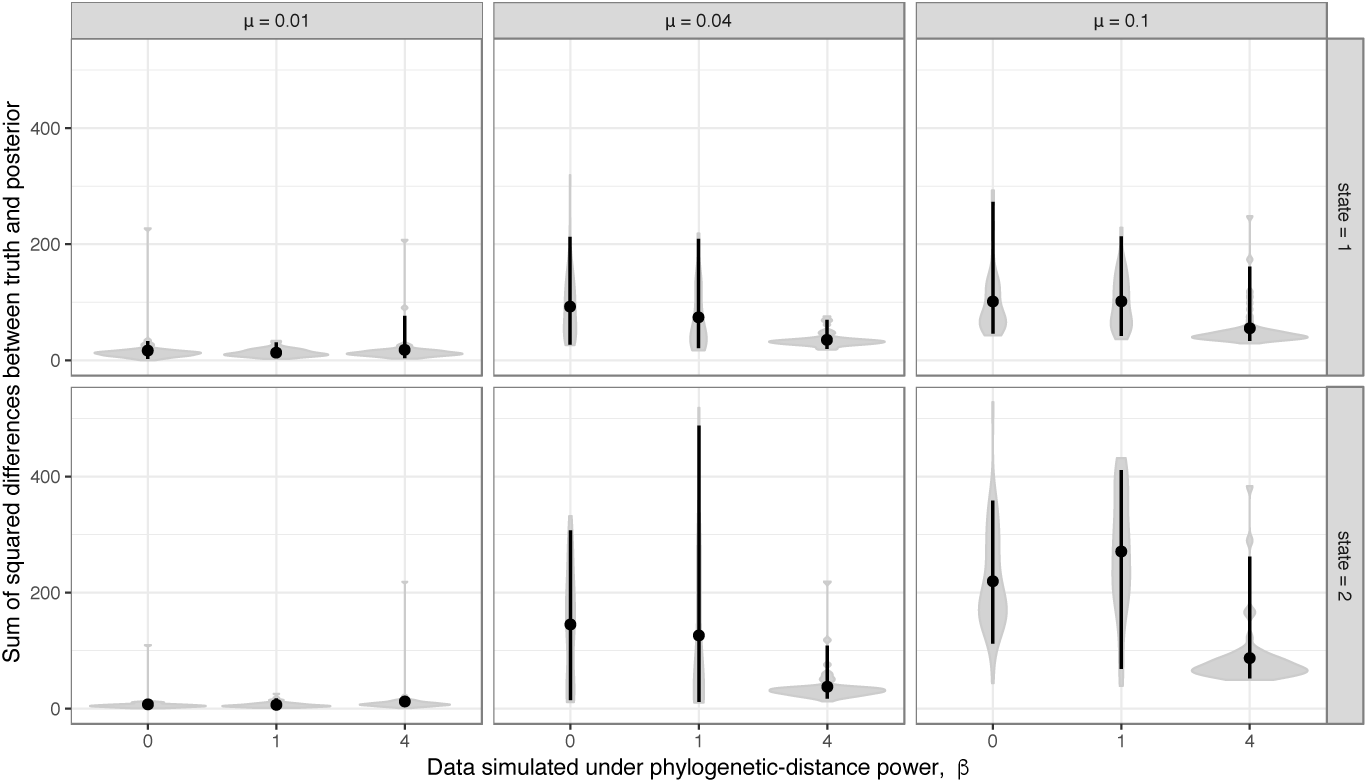
Errors for inferred dispersal histories of simulation study. The sum of squared differences between the posterior probability (0 ≤ P ≤ 1) and the true history (P = 0 or 1) for neach host and each internal node were computed per simulated dataset. Each violin plot shows the distribution of these sums for each batch of 50 simulated datasets. Means are represented by black dots, black vertical lines show the 95% CI. Values of phylogenetic-distance power (*β*) are shown in the x-axis, columns are separated by the host-repertoire evolution rate (*µ*), and each row shows the error on the inference of each character state, i.e. potential host (1) or actual host (2).

**Figure 6:**
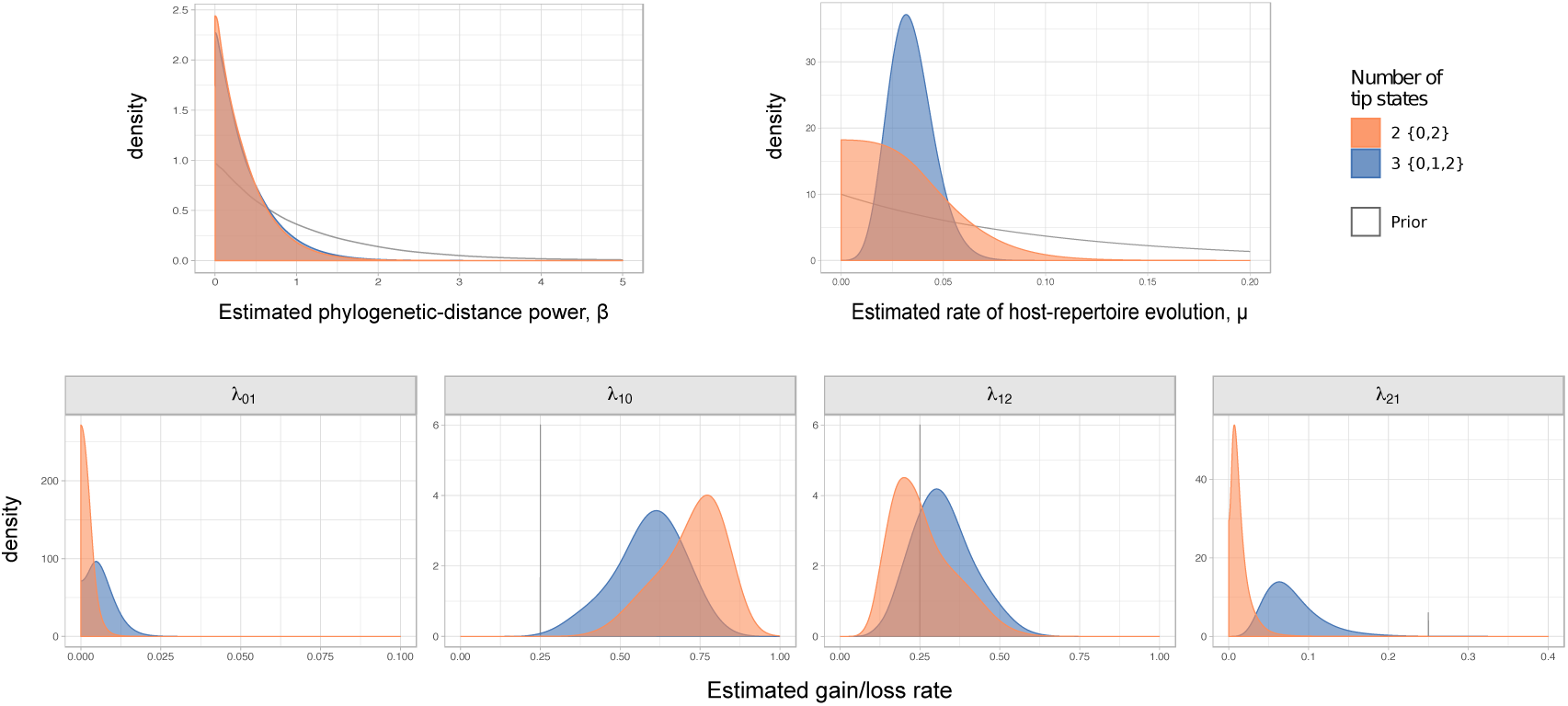
Marginal posterior densities for parameters in the Nymphalini-Angiosperms study for both the full dataset (3 states at tips) and the dataset omitting the intermediate state (2 states at tips). Grey lines corresponds to the priors *β* ∼ Exponential(1), *µ* ∼ Exponential(10), and *λ ∼* Dirichlet(1, 1, 1, 1).

**Figure 7:**
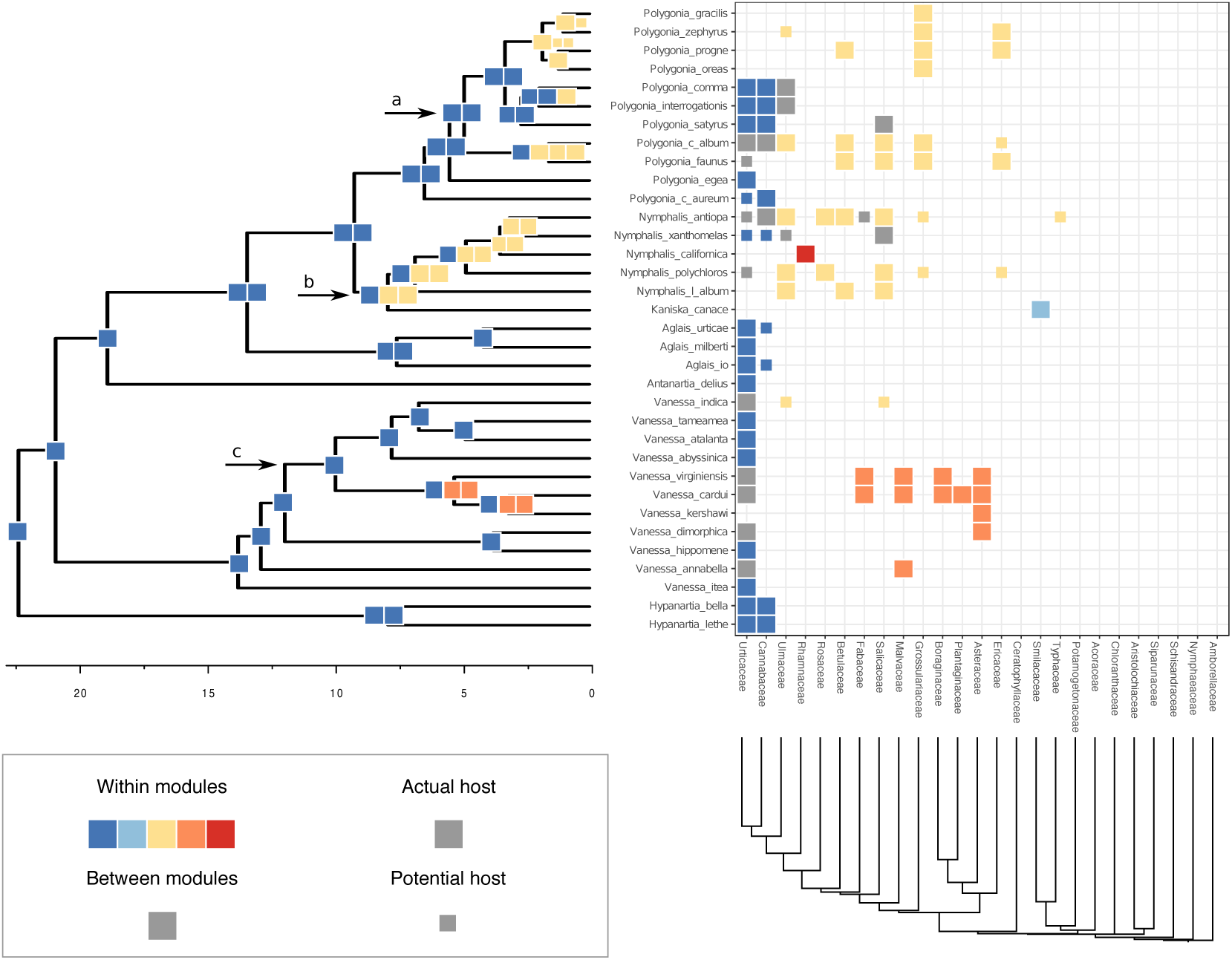
Evolution of butterfly-plant interactions through time. Ancestral state estimates (left) of host repertoire across the Nymphalini phylogeny are shown for interactions with more than 75% posterior probability. The x-axis under shows time before present in millions of years. Extant species interactions (right) between Nymphalini and their host plants are presented as a raster, where each square represents one interaction between a butterfly species and a host family. Colors represent different modules, i.e. groups of plants that are often hosts to the same butterflies at present time. Square size was used to differentiate between actual and potential hosts. Arrows indicate nodes shown in Fig. 8.

Importantly, the empirical analysis indicates that the method is able to model the evolution of fundamental and realized host repertoires even when the information about potential hosts is lacking. This significantly increases the applicability of our method, as information about fundamental host repertoires is missing for most host-parasite systems. Potential host data is difficult to collect, as it requires experimental testing of a large number of potential host-parasite pairs. A possible improvement of our method, which we did not explore here, would be to model uncertainty in the observations of non-hosts when data on potential hosts are missing. That is, if we had no information about a host species being used by a particular parasite, we would translate that to a certain probability *p* of the species actually being a non-host, and a complementary probability 1 *– p* of it being a potential host (Kuhner and McGill 2014). Modeling this observational uncertainty could help reduce the bias in parameter estimates that we observed when data on potential hosts were missing and all 0 states in the dataset were inappropriately treated as true non-hosts. This extension would also allow us to make predictions about host use abilities in extant parasites. These predictions could then inform experiments that aim to characterize fundamental host repertoires.

We demonstrated the empirical application of our approach with a Bayesian inference of the coevolutionary history between 34 Nymphalini butterflies and 25 angiosperm families. We estimated the rate of host repertoire evolution along each branch of the butterfly tree as being between 0.33 and 0.93 events per million years. Bayes factors favored the independence model, where the probability of gaining a given hosts is not affected by the phylogenetic distance between hosts. As explained above, this does not necessarily mean that host relatedness plays no role, only that the effect is not large enough for us to detect it with the current approach and the given data.

Estimates of gain and loss rates were not symmetric, and the rates also varied between states. According to our results, gain of the ability to use a host, *λ*_01_, is very rare (0.5% to 1.9% of overall rate), whereas loss is common (47% to 73% of overall rate). On the other hand, transition rates between states 1 and 2 were more symmetric and gain is more common than loss (*λ*_12_ between 15% and 39%; *λ*_21_ between 6% and 18% of overall rate). These rate estimates support the idea that the use of the same host lineage by multiple, phylogenetically widespread butterfly lineages is more likely explained by recolonization of hosts that have been used in the past (recurrence homoplasy), that is, by transitions between actual and potential hosts, rather than by completely independent colonizations of the same host (Janz et al. 2001). Note that alternative scenarios that have been proposed in the literature to explain the evolution of Nymphalini host plant preferences, for instance by involving narrow ancestral host plant ranges and repeated independent colonization events, are also allowed by our model, but they are inferred to be much less likely than the conservative host repertoire scenario. Yet, because the potential host state is exited at the highest rate, the rate estimates also suggest that parasites do not retain their potential host relationships for prolonged periods of time. The moderate rates of transitions between potential and actual host states and the high departure rate from the potential host state together help explain why phylogenetic “pulses” of recurrent host acquisition manifest in some lineages but not others.

For example, the use of Grossulariaceae by two non-sister clades within *Polygonia* is best explained by a scenario where Grossulariaceae was a potential host for the ancestral species (node 60 in Fig. 8) and was subsequently gained as an actual host twice (at nodes 53 and 58, Fig. S4). The ability to use Salicaceae host plants seems to be even older. Salicaceae was likely a potential host for the ancestor of *Nymphalis* + *Polygonia* and later became an actual host in three different parts of the clade. If potential hosts were not explicitly modeled here, these transitions would look like three independent colonizations of a plant group that is very distant from the ancestral host (Salicaceae and Urticaceae diverged about 90 Ma). Instead, we could show that what might appear as big and sudden host shifts, are in fact the result of retention of ancestral host use abilities.

**Figure 8:**
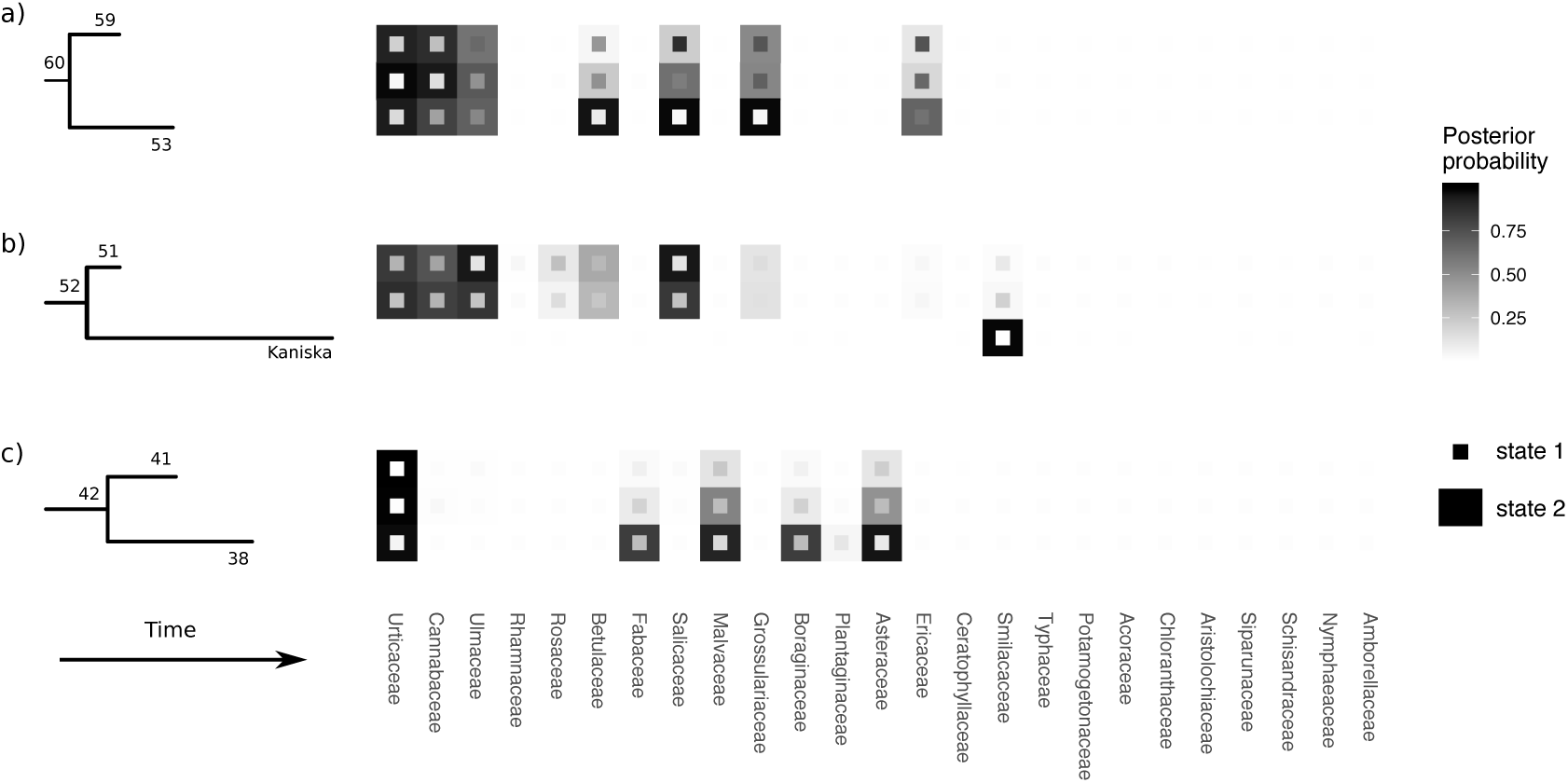
Host repertoires at selected nodes of the Nymphalini tree (arrows in Fig. 7). Numbers indicate the node index (compatible with Fig. S3). For the only terminal taxa depicted, *Kaniska canace*, the observed host repertoire is shown. For all other repertoires, the posterior probabilities for states 1 and 2 are shown.

Understanding how ecological interactions change is crucial if we want to predict both short and long-term consequences of global mixing of biota (Hoberg and Brooks 2015). Host-parasite interactions are of particular interest given the risk of emerging diseases, which can affect human populations directly and indirectly through their effects on crop species and wildlife (Brooks et al. 2014). Our method was designed to quantify changes in host-parasite associations by modeling the process of gaining and losing hosts, thus allowing us to make predictions based on host-parasite history. Hopefully, our approach will not only generate deeper insights into the evolutionary dynamics of host-parasite associations but also help humankind mitigate some of the risks incurred by current environmental change.

## Funding

M.J.L. was supported by the Donnelley Fellowship through the Yale Institute of Biospheric Studies, with early work in this study supported by the NSF Postdoctoral Fellowship in Biology (DBI-1612153). The Swedish Research Council supported SN (2015-04218) and FR (2014-05901).

*

## Notes

https://github.com/mpiresbr/host_repertoire

