## Supplementary Information for "Bayesian inference of ancestral host-parasite interactions under a phylogenetic model of host repertoire evolution"

**References for host use records for Nymphalini.**

**Figure S1: Empirical dataset used to validate the model.**

**Figure S2: Modules of Nymphalini-angiosperms interactions.**

**Figure S3: Time-calibrated phylogeny of Nymphalini.**

**Figure S4: Ancestral state reconstruction of Nymphalini host repertoire.**

**References used to compile host records for Nymphalini:**


Braby MF (2000) Butterflies of Australia: Their identification, biology and distribution. Vol 2. CSIRO Publishing, Collingwood.

DeVries PJ (1987) The Butterflies of Costa Rica and their Natural History, Papilionidae, Pieridae, Nymphalidae. Princeton University Press, Princeton.

Ebert G (1993) Die Schmetterlinge Baden-Wrttembergs. Verlag Eugen Ulmer, Stuttgart.

James DG and Nunnallee D (2011) Life histories of Cascadia butterflies. Oregon State University Press, Corvallis, OR.

Janz N, Nyblom K and Nylin S (2001) Evolutionary dynamics of host-plant specialization: A case study of the tribe Nymphalini. Evolution 55: 783-796.

Larsen TB (1991) The butterflies of Kenya and their natural history. Oxford University Press, Oxford.

Larsen TB (2005) Butterflies of West Africa. Apollo Books, Stenstrup.

Layberry RA, Hall PW and Lafontaine JD (1998) The butterflies of Canada. University of Toronto Press, Toronto.

Migdoll I (1987) Field guide to the butterflies of Southern Africa. C. Struik, Cape Town.

Savela M (2014) Lepidoptera and some other life forms: http://www.nic.funet.fi/pub/sci/bio/life/intro.html

Scott JA (1986) The butterflies of North America. Stanford University Press, Stanford, CA.

Tennent J (1996) The butterflies of Morocco, Algeria and Tunisia. Gem Publishing Company, Wallingford, Oxfordshire.

Tolman T and Lewington R (1997) Collins field guide: Butterflies of Britain and Europe. Harper Collins Publishers Ltd., London.

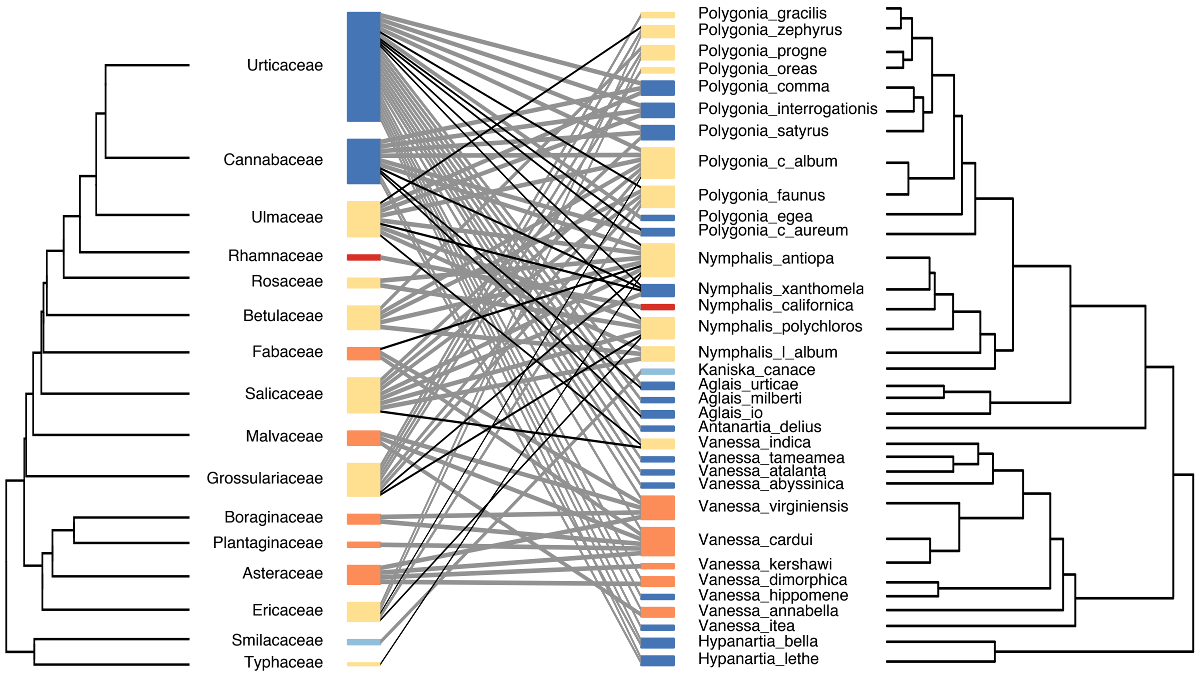


Figure 1: Empirical dataset used to validate the model. Interactions between Nymphalini butterflies (right) and angiosperm families (left) with taxa ordered by phylogenetic relationship. Thickness of boxes is proportional to the number of interactions. Box colors show the five modules found in the dataset (consistent with figs. 7 and S2. Grey lines represent interactions with actual hosts and black lines represent interactions with potential hosts.

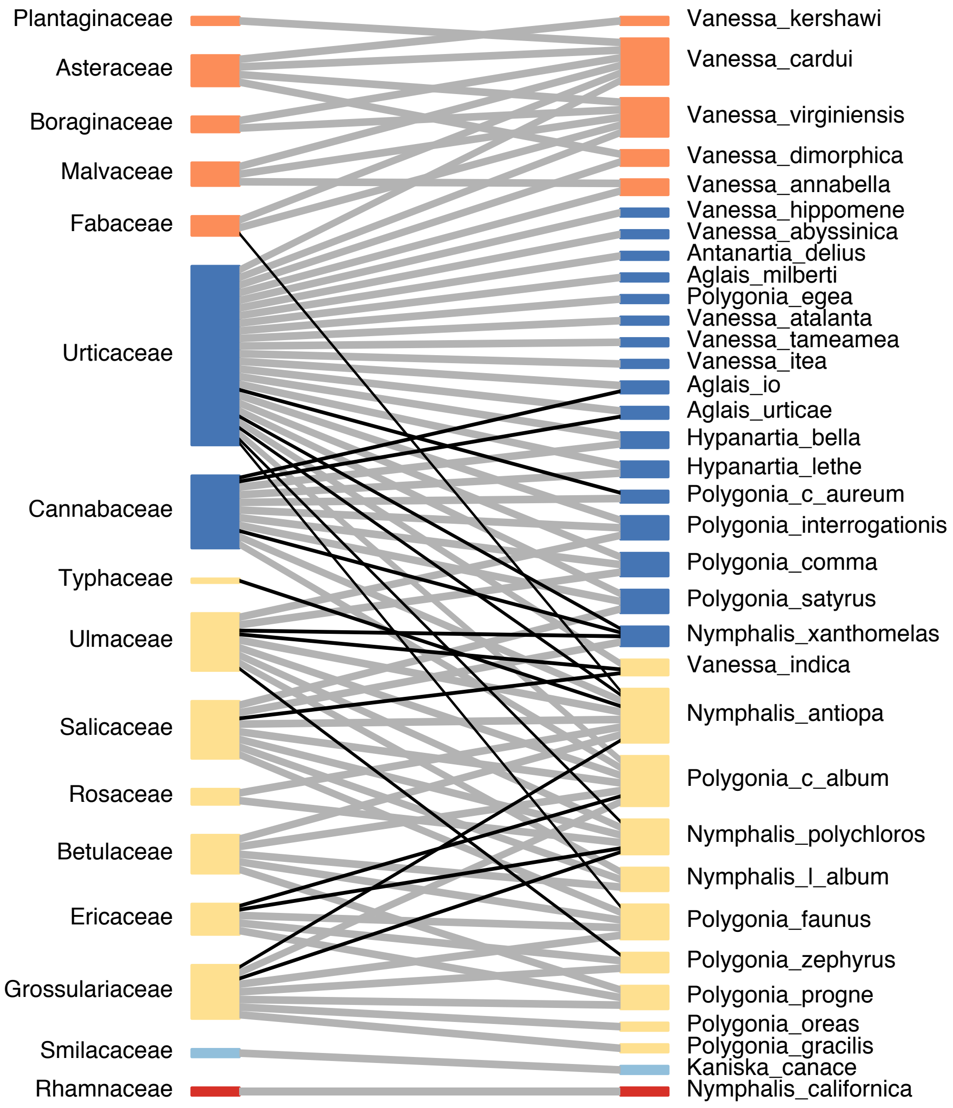


Figure 2: Interactions between Nymphalini butterflies (right) and angiosperm families (left) with taxa ordered to show the five modules identified in this dataset (colors). Grey lines represent interactions with actual hosts and black lines represent interactions with potential hosts.

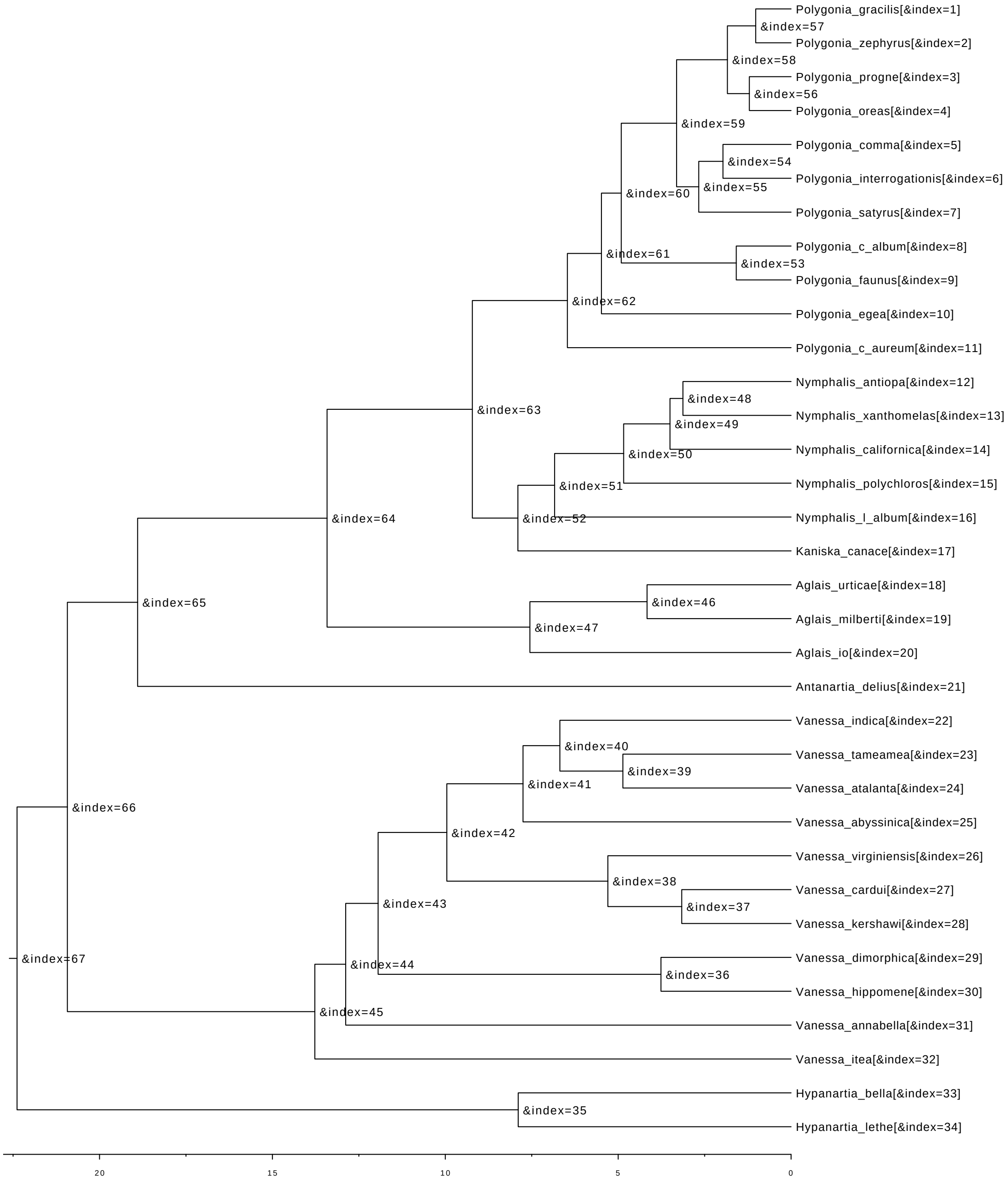


Figure 3: Time-calibrated phylogeny of Nymphalini tribe from Chazot et al. (unpublished). Indices of internal nodes correspond to row names in Figure S4.

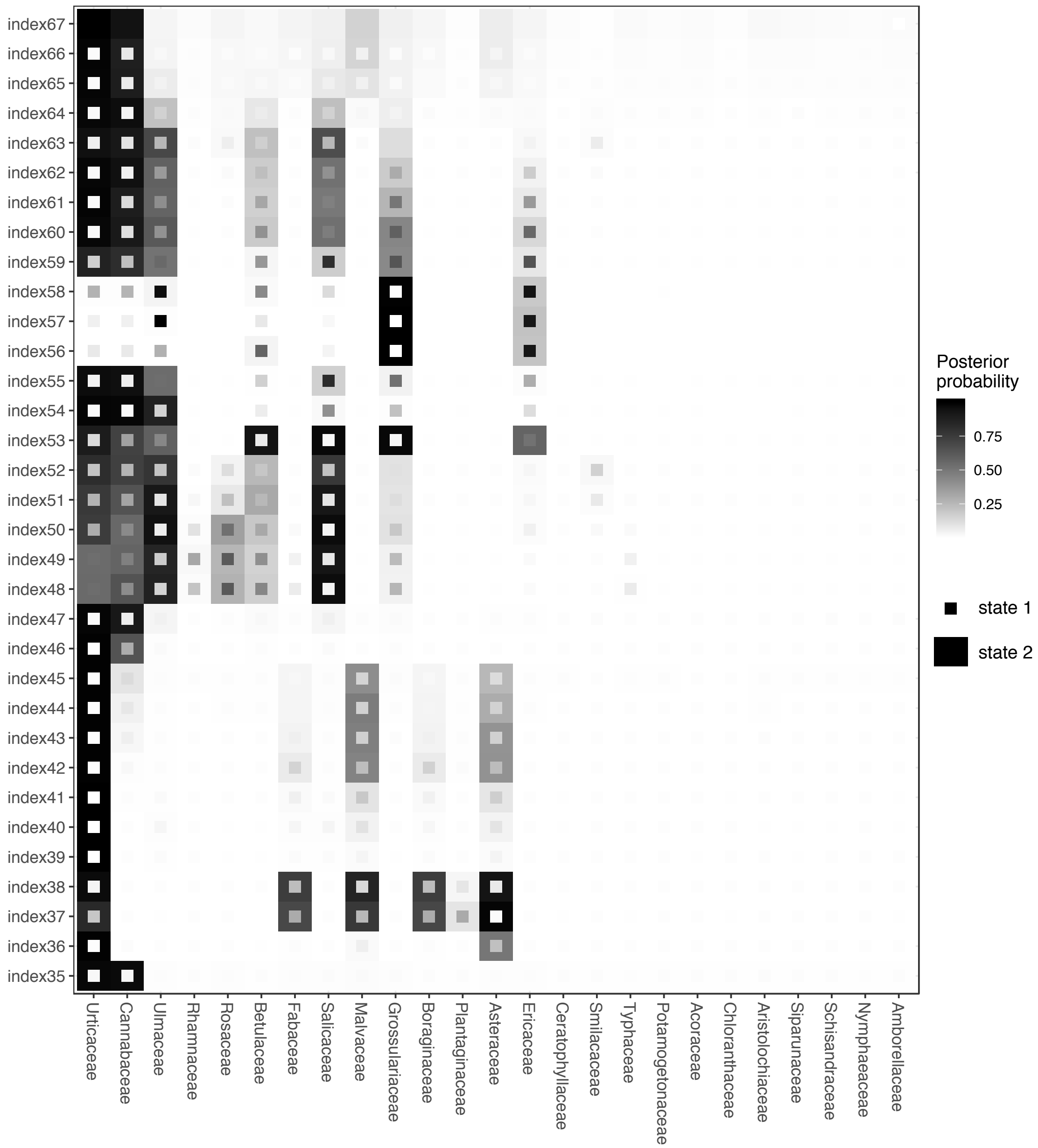


Figure 4: Posterior probability for ancestral host repertoires of Nymphalini butterflies. Rows show the host repertoires at internal nodes of the Nymphalini phylogeny (same index as in Fig. S3). Small squares represent potential hosts (state 1), whereas large squares represent actual hosts (state 2). The probability of a host being on state 1 and on state 2 is shown by the color scale.
